# Sick Bats Stay Home Alone: Social distancing during the acute phase response in Egyptian fruit bats (*Rousettus aegyptiacus*)

**DOI:** 10.1101/2020.07.06.189357

**Authors:** Kelsey R. Moreno, Maya Weinberg, Lee Harten, Valeria B. Salinas Ramos, L. Gerardo Herrera M., Gábor Á. Czirják, Yossi Yovel

## Abstract

Along with its many advantages, social roosting imposes a major risk of pathogen transmission. How social animals, and especially free-ranging mammals, reduce this risk is poorly documented. We used lipopolysaccharide injection to imitate bacterial sickness in both a captive and a free-ranging colony of an extremely social, long lived mammal – the Egyptian fruit bat. We monitored behavioral and physiological responses using an arsenal of methods, including on-board GPS and acceleration, video, temperature and weight measurements, and blood samples. Sick-like bats exhibited an increased immune response, as well as classical illness symptoms including fever, weight loss, anorexia, and lethargy. Notably, they also isolated themselves from the group by leaving the social cluster and avoiding contact. Free-ranging individuals ceased foraging outdoors for at least two nights. Together, these sickness behaviors demonstrate a strong, integrative immune response which promotes recovery of infected individuals while protecting their group members from transmission of pathogens, and at the same time, reducing spillover events outside the roost.

## 1. Introduction

Sickness behavior, as first described by Hart^1^ is a set of behavioral changes that ill individuals develop simultaneously with their illness including lethargy, depression, anxiety, malaise, loss of appetite, sleepiness, hyperalgesia, reduction in grooming and general movement, and a loss of interest in their surroundings. These behaviors are well preserved across vertebrates and have been documented in some invertebrates^2,3^. Although first believed to be a side effect of immunological processes, sickness behavior is now agreed to be an adaptive behavioral trait that supports the physiological struggle of a sick individual against the source of infection^4^. Sickness behavior has also been suggested as a mechanism to reduce transmission of pathogens to kin^5^ and within the social group^6^, a feature that might be critical for animals living in dense social populations. A few documented examples include social isolation of sick individuals among eusocial insects such as bees^7,8^ and ants^9–12^. So far, social isolation in vertebrates has been shown to be a case of avoidance by healthy individuals. In multiple vertebrate species, healthy individuals can discriminate between sick and healthy conspecifics and spend more time in proximity to healthy than sick individuals^13–15^. Additionally, there is mounting evidence that sickness does not automatically induce classical sickness behaviors like lethargy, but may be exhibited to varying degrees in species with different life history strategies^16^ or can be suppressed during social contexts as seen in male zebra finches^17^. Such variation demands refinement of the classical sickness behavior hypothesis to account for interspecific and contextual variation. Importantly, sickness behavior has been rarely investigated in free-ranging animals^18,19^ with only two studies on free-ranging wild mammals demonstrating behavioral changes during sickness^20,21^.

Current knowledge of the metabolic and systemic changes during the first few days post injury or immunological challenge (termed the acute phase response^22^) in bats suggests that different species might respond differently to the same threat. Immune responses have been measured in a few bat species using lipopolysaccharide (LPS), a bacterial endotoxin that induces an inflammatory response by stimulating the release of cytokines^23,24^. Fever has been documented in response to an LPS challenge in three out of four species examined (*Myotis vivesi*^25^, *Carollia perspicillata*^26^, *Artibeus lituratus*^27^, but not in *Molossus molossus*^28^). Some species exhibit clear leukocytosis (*Carollia perspicillata*^29^ and *Desmondus rotundus*^30^), while others shift the ratio of white blood cell types without a clear overall increase (*Artibeus literatus*^27^) and others do not increase white blood cell production at all (*Molossus molossus*^28^). Weight loss was observed in all instances body mass was measured, likely due to increased metabolic rate, as documented in the great fruit-eating bat^27^ and fish-eating bat^25^, and decreased food consumption, as observed widely across animals^3^ and recently documented in bats^31^.

Many bats are extremely social, roosting in large colonies and clustering in tight groups. Still, sickness behavior and its role in preventing pathogen transmission is poorly studied in bats. In the only previously studied species, LPS-injected vampire bats (*Desmodus rotundus*) decreased overall activity levels, reduced grooming towards others, received less grooming from group mates^30,32^, and had lower network centrality^21^ yet there was no reduction in the number of food donors or the amount of food received following a night of food deprivation^32^. Such behavioral changes support the hypothesis that sickness behaviors alter interactions by which diseases are transmitted^5,30^.

Molecular aspects of the immune response are also understudied in bats. Haptoglobin is a major acute phase protein with bacteriostatic and immunomodulatory effects^33^. In bats, haptoglobin levels increased in response to an immune challenge with a fungal antigen^34^. Lysozyme is a highly conserved antibacterial enzyme^35^, which hydrolyzes cell wall peptidoglycan and modulates the immune response to infection^36^. Despite its key role in the immune response, lysozyme has only been recorded in bats as a component of the digestive system^37^ and as a measure of environmental disturbance^38^.

In this study we assess the acute phase response (APR) and sickness behavior of the highly social Egyptian fruit bat (*Rousettus aegyptiacus*), in captive adult and free-ranging juvenile individuals, using LPS to simulate an infection without use of an infectious pathogen. We hypothesized that Egyptian fruit bats would follow general mammalian sickness physiology, though due to inter-specific variation in bat sickness responses, we could not predict the severity or details of the response. Behaviorally, we expected a reduction in movement and food consumption, and avoidance of sick-like individuals by healthy individuals. Our findings demonstrated that physiologically, Egyptian fruit bats have responses between those of other bats, raising body temperature and reducing food intake, but not exhibiting leukocytosis. Behaviorally, they responded very strongly. Unlike previous reports in other mammals, sick-like individuals abandoned the social cluster and remained distant from conspecifics. This reaction was observed in both captive and free-ranging individuals. GPS tracking demonstrated that free-ranging sick-like individuals failed to leave the roost to forage. We suggest that the inflammatory reaction and the sickness behavior together serve to conserve resources while maximizing swift bacterial eradication and reducing transmission among group members.

## 2. Results

We established two closed colonies from a mixture of 19 recently caught male and 18 previously housed female Egyptian fruit bats (Rousettus aegyptiacus). Each colony contained 5 bats which were challenged with a lipopolysaccharide (LPS) injection, 5 bats which were injected with phosphate buffered saline (PBS) buffer as a control, and additional bats to provide a full social environment. Following these trials, we challenged 5 bats in an open colony where bats have free access to nature at all times39. These individuals also received an injection of PBS at a separate time, and thus served as their own controls. Data were collected pre and post injection using a combination of on-board trackers, infrared video cameras, body weight measurements, and blood draws.

### a. Evidence of Illness Response

The immune-challenged bats showed a clear physiological sickness-like response. Skin temperature of the challenged group in the closed colony rapidly elevated following injection (Figure 1 a; p = 0.003 main effect of time, p = 0.02 interaction between time and treatment group; Mixed ANOVA with temperature as the response variable, treatment as a between subjects predictor, and time as a within subjects predictor). Skin temperature reached its peak at around 80 minutes after injection at 37.6 ± 1.3 °C (Mean±SD). Skin temperature was elevated continuously for up to 36 hours post challenge, though only in the first 16-22 hours was there a stark contrast between challenged (37.0±1.13 °C) and control means (35.7±1.5 °C). We observed clear spikes in skin temperature in both the control and challenged groups every time the bats were removed from the colony room for weighing and taking blood samples. We suspect this is due to stress associated with the handling process, as stress has been demonstrated to cause body temperature elevation in bats40. We did not measure the temperature of the free-ranging bats.

**Figure 1.**
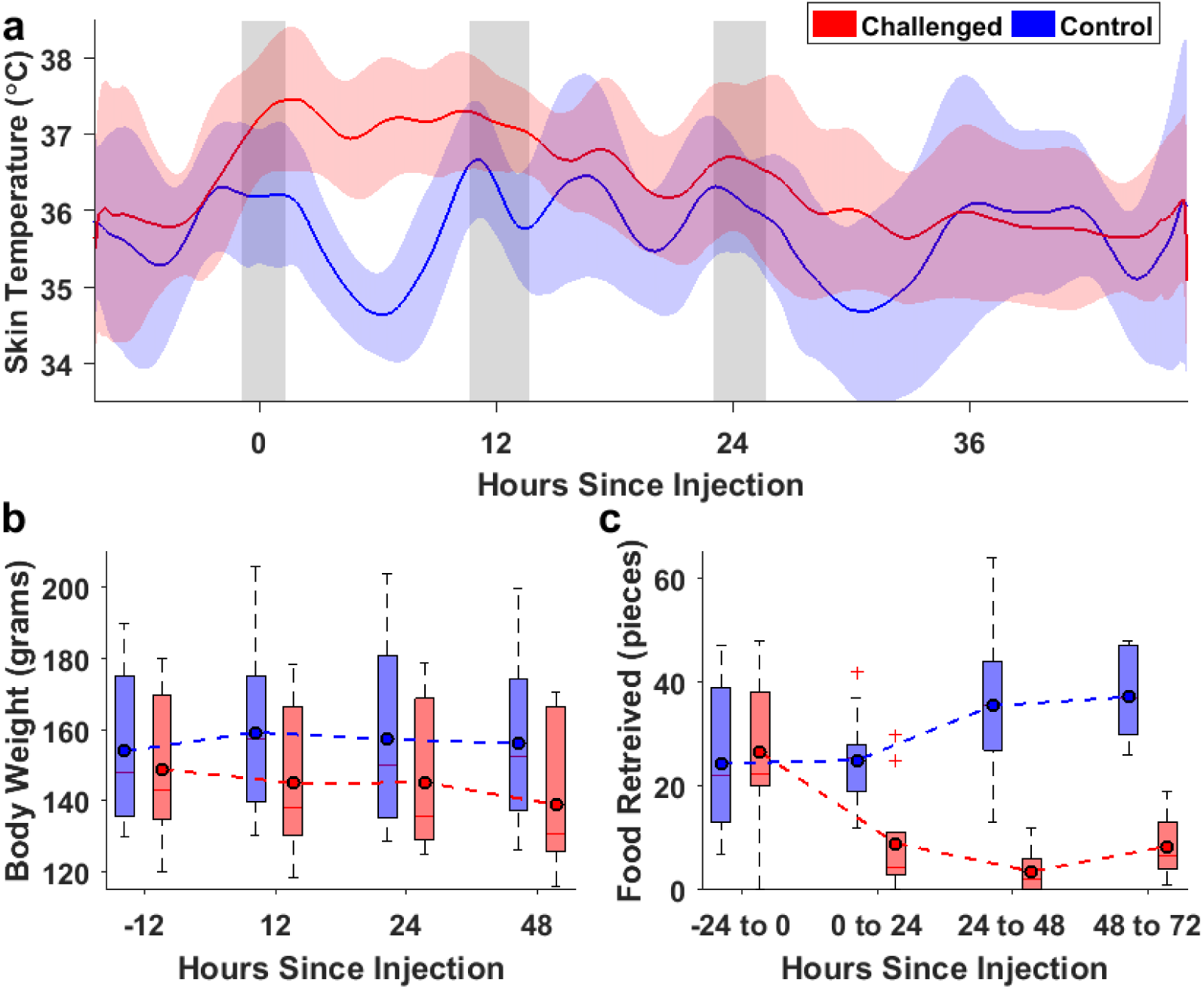
Evidence of illness response in LPS-challenged bats. (a) Skin temperature of Challenged bats (n=5) was higher than controls (n=6), mostly in the first 12 hours. Shading for each group’s respective mean line shows the 95% confidence interval. Grey shadowed areas depict handling periods, which probably led to a temperature elevation. (b) Challenged bats (n=10) lost weight while controls (n=10) did not. (c) Challenged (n=10) bats exhibited anorexia.

In the closed colony, only sick-like bats lost weight, losing on average 9.9 ±4.5g over 48 hours (Figure 1 b; p < 0.001 interaction between time and treatment; Mixed ANOVA as above, with body weight as the response variable). Moreover, monitoring the food bowl showed that weight loss was at least partially a result of anorexia, as individuals markedly reduced food consumption following LPS injection (Figure 1 c; p < 0.001 main effect of treatment, p = 0.044 main effect of time, p < 0.001 interaction; Mixed ANOVA as above, with pieces of food retrieved as the response variable). Bats in the open colony also lost weight, but the degree of loss was only nearly-significant, likely due to small sample size and the age of these individuals (p = 0.064; paired t-test with weight as the response variable and time as a within subjects predictor).

### b. Bats isolate themselves when they are sick

Distance to nearest neighbor (mm) and isolation level (categorical 0-3) in the closed and open colonies, respectively, was significantly explained by the time of day, treatment group, and whether it was pre or post LPS injection (Figure 2 a, b; p < 0.01 for all main effects and interactions; Mixed effect GLMM with treatment group, time of day, and pre/post LPS injection as fixed effects and bat ID set as random effect). Healthy individuals in both colonies displayed a cyclical pattern of distance from their nearest neighbor following natural circadian rhythms, with the lowest distances occurring during the daytime sleep when the bats tightly clustered together and the highest distances during the nighttime when bats were active. Following LPS injection, sick-like bats in both the closed and open colonies deviated from this pattern and increased distances from the bat cluster during the daytime (Figure 2 c). Interestingly, sick-like bats isolated themselves from the cluster instead of being rejected by other group members. Video observations revealed that they either withdrew from the bat cluster or did not join the cluster upon its formation (Supplementary Videos: Video1, Video2, Video3, Video4, Video5). This self-isolation behavior lasted approximately two days. Moreover, during this isolation period, sick-like individuals displayed dramatically reduced movement, as we quantified with accelerometers attached to the animals in the closed colony (Figure 2 d, e; p = 0.036 effect of treatment, p < 0.001 effect of time of day, and p = 0.002 interaction between time and treatment; Mixed effect GLMM as above, with proportion of time in high activity as the response variable). The action of moving out of the cluster for self-isolation was a behavioral exception in light of their general tendency to remain still.

**Figure 2.**
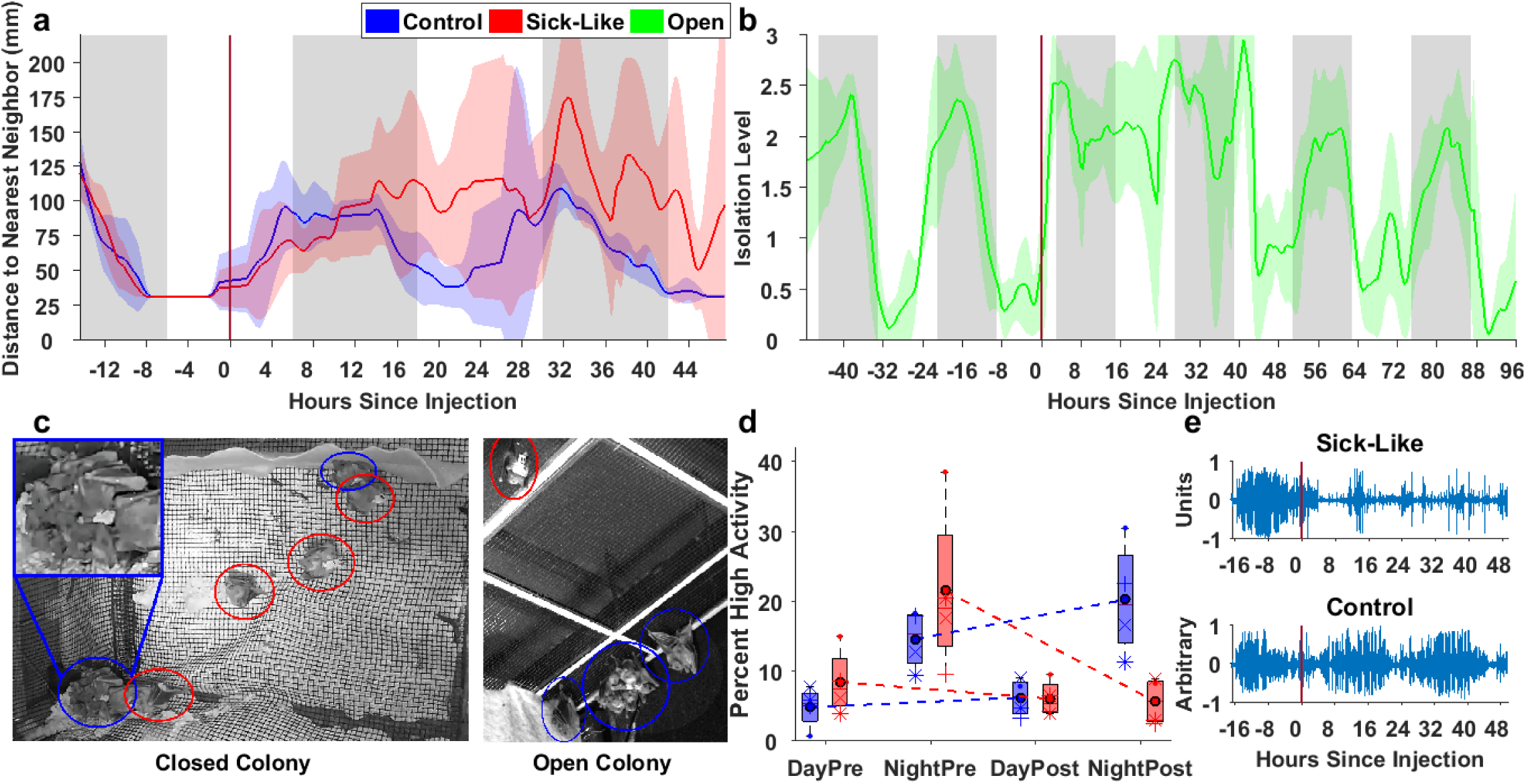
Sick-like bats exhibit social isolation. Sick-like bats in both the closed (a; n=5) and open (b; n=5) colonies maintained larger distances from nearest neighbor. Nights are indicated by grey vertical shading. Confidence intervals (95%) are shown through shading for each group’s respective line representing the mean. (c) Photos from the closed (left) and open (right) colony show sick-like individuals (circled in red) out of the bat cluster (shown in greater detail in the top left inset) and distant from conspecifics (circled in blue). Note the single control individual in the closed colony (top right) that chose to perch near sick-like individuals (see Discussion). (d) Challenged (n=4) bats moved much less than control bats (n=4). (e) This is also shown with an acceleration sequence example for the full duration of the study (approximately 64 hours) for a challenged bat (top) and a control bat (bottom) with the moment of the treatment marked by a vertical dark red line.

### c. Bats Stay in When Sick

Sick-like bats in the open colony dramatically changed their foraging behavior. Following the LPS injection, individuals stayed in the colony for at least 2 nights and up to 5 nights, whereas before the injection they consistently exited to forage (Figure 2a; p < 0.001; Fisher’s exact comparing the probability of exiting before and after injection). Moreover, once individuals resumed foraging, they initially flew shorter distances than they had prior to LPS injection (Figure 3 b, c, Supplementary figure 1; p = 0.004; Repeated ANOVA with distance flown as the response variable and time as a within subjects predictor). We validated that the staying in behavior was not an artifact of the food supplement that was given in the open colony by showing that these bats ate significantly less than expected given their body weight (Supplementary figure 2; p = 0.012; post-hoc: p = 0.012 night 1 to expected, p = 0.044 night 2 to expected; Repeated ANOVA as above with pieces of food retrieved as the response variable). The decrease in food consumption was similar in the two colonies (bats consumed 23.39±24.83% vs. 49.87±29.6% of the expected amount in the closed and open colonies, respectively).

**Figure 3.**
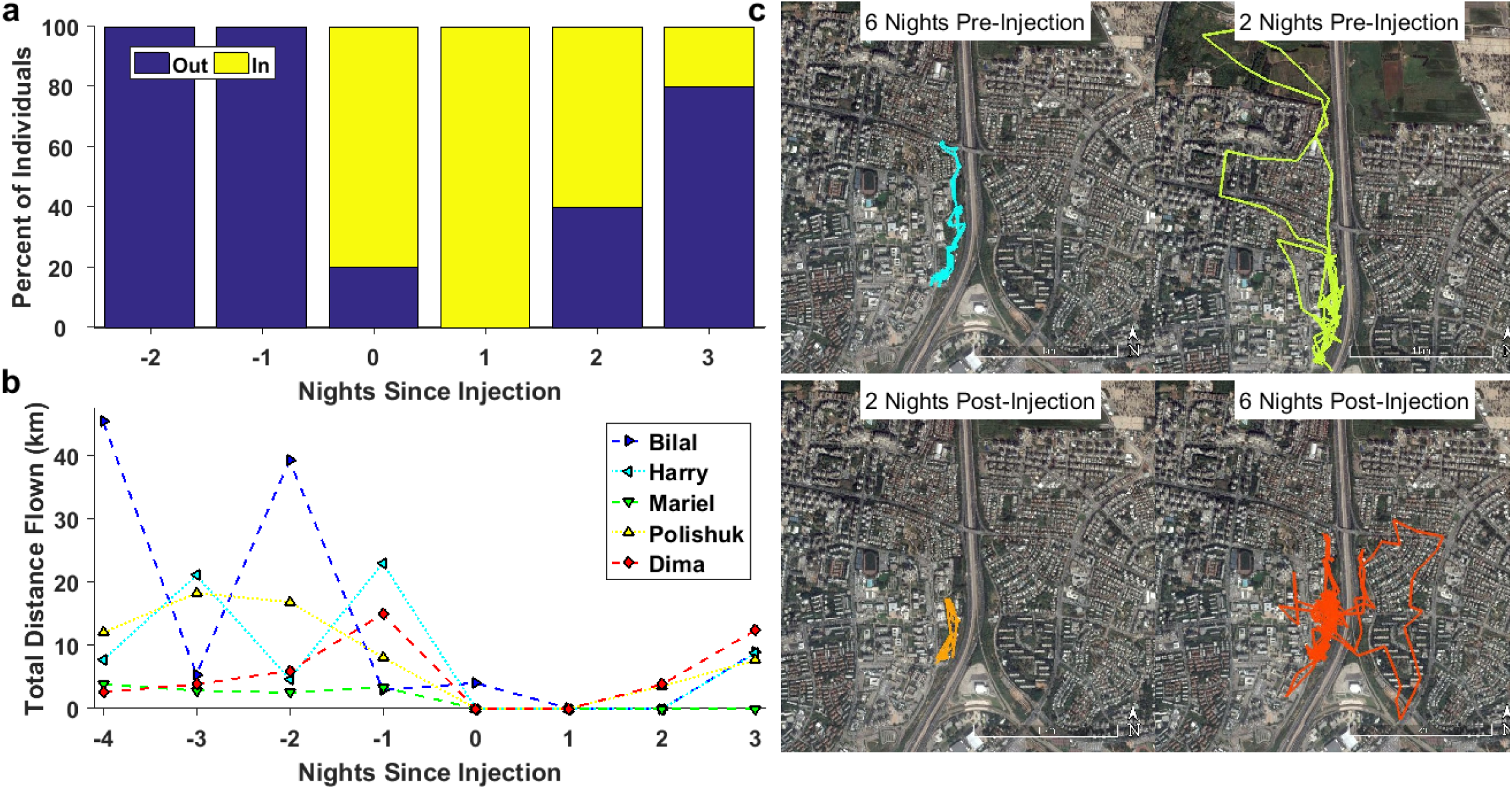
Sick-like bats stop foraging. Following LPS injection, individuals (n=5) in the open colony (a) did not exit to forage, as shown in yellow, and (b) flew shorter distances on their first trip back out than on trips prior to injection. This can also be seen through GPS tracks of one individual (Polishuk) with examples (c) prior to injection (top, in blue and yellow), on the first night out after injection (bottom left, in orange), and after recovery (bottom right, in dark orange).

### d. Acute Phase Response in Fruit Bats

Immunologically, there was a clear acute phase response as demonstrated by multiple blood parameters. Total white blood cell count showed no significant difference between challenged and control groups at all times in both the closed and open colonies. However, the neutrophil to lymphocyte ratio (NLR), a ratio of the count of the two most involved white blood cell types in the acute phase response that also serves as an indicator of physiological stress, was found to be significantly higher in the treatment group following LPS injection for the closed colony (Figure 4a; p = 0.007 interaction between treatment and time; Mixed ANOVA as above, with NLR as the response variable). In the closed colony, the challenged group maintained a similar NLR throughout the experiment, while the control group’s ratio decreased after the initial measurement. The NLR was higher prior to LPS injection in the open colony (p = 0.015; Paired t-test as above with NLR as the response variable).

**Figure 4.**
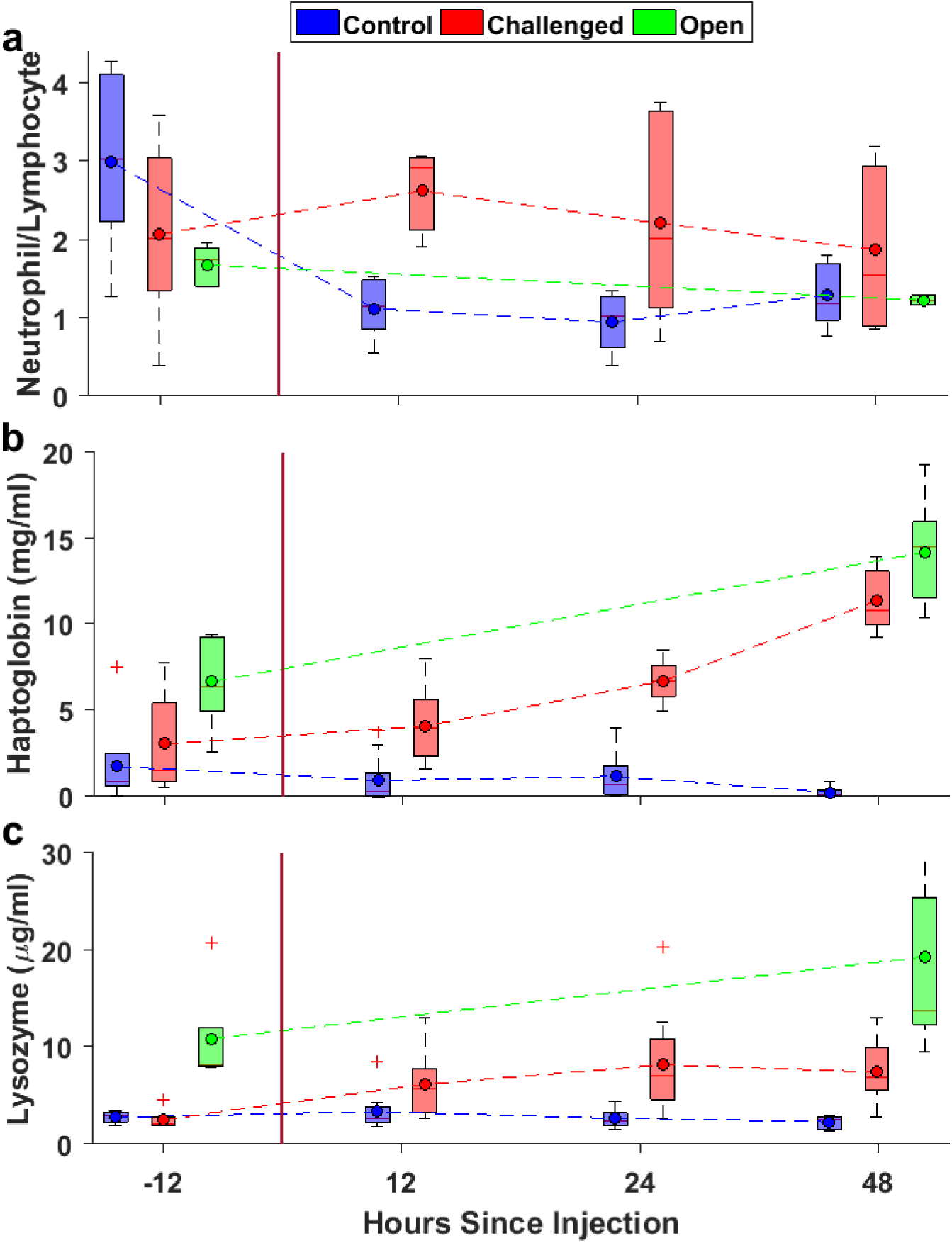
Immunological responses in both the closed (red – challenged, n=10; and blue – control, n=10) and open (light green – n=5) colonies are shown for the (a) neutrophil to lymphocyte count ratio, (b) haptoglobin concentration, and (c) lysozyme concentration.

The levels of haptoglobin were significantly higher following LPS injection (Figure 4b). In the closed colony, haptoglobin increased over time (p = 0.001 interaction between treatment and time; Mixed ANOVA as above, with haptoglobin as the response variable;) and was significantly higher after injection in the challenged group than the control (p < 0.001), particularly after 24 hours (p = 0.004) and 48 hours (p < 0.001). In the open colony, the pups had higher haptoglobin levels after 48 hours than prior to LPS injection (p = 0.006; Paired t-test as above with NLR as the response variable). The levels of lysozyme were higher following LPS injection in the experimental group in the closed colony but not in the open colony (Figure 4c; p = 0.033 interaction between treatment and time; Mixed ANOVA as above, with lysozyme as the response variable; p = 0.25; Paired t-test as above with NLR as the response variable; respectively).

## 3. Discussion

Egyptian fruit bats present a unique opportunity to understand sickness responses in long lived, highly social mammals that live in densely populated, high-contact groups. We sought to understand the behavioral and physiological response to bacteria, a common yet relatively understudied cause of illness, in *R. aegyptiacus*. We used a standard immune challenge in order to induce an acute phase reaction by the innate immune system and the accompanying sickness behavior. Despite using a relatively medium dosage of LPS compared to similar experiments in other bats species^27,29,30,32^, our bats were very clearly sick. LPS-challenged bats had fever, lost weight, consumed less food than before the challenge, and developed clear sickness behavior, including lethargy, reduction in social interaction, and general movement.

Skin temperature was found to be a reliable proxy of body temperature, as the values measured pre-challenge are known to fit healthy normal fruit bats^41^. Body temperature elevation in response to an immune challenge is observed across bat species, as it is in other mammals, although Melhado et al^31^ point out this only occurs in bats challenged during the resting phase^25–27,29^, and not for bats challenged during the active phase^28,31^. We challenged bats in both active and resting phases, though we did not have a large enough sample to compare between the two. The observed temperature elevation in our study of 1 to 1.5 °C is similar to that found in other bat species including *C. perspicillata*^29^ and *A. lituratus*^27^. Interestingly, *Rousettus* elevated fever was maintained for approximately 36 hours post challenge, much longer than reported for other bat species (up to 6 hours).

During sickness, bats isolate themselves and hang apart from conspecifics either by failing to join a cluster during its formation or by leaving the cluster during the daily sleeping hours. This behavior was initiated spontaneously by the sick-like bat and was not a response to any behavior of the healthy controls. Sick-like individuals then maintained their isolation for about two days. While general listlessness during illness clearly contributes to distance from conspecifics, instances of individuals leaving the cluster or changing locations within the room from one isolated location to another demonstrate this is not the sole explanation. To our knowledge, this is the first documented instance detailing this self-isolation behavior in a mammal. Such self-isolation behavior is notably different from what has been observed in other non-human mammals. Despite social withdrawal being considered a hallmark of sickness behavior^1^, previous findings of mammalian social isolation are of healthy individuals avoiding sick conspecifics^13,42,43^, or sick individuals reducing interactions^32,44^ or shared space use^17^ due to lethargy. Moving outside of the bat cluster might also come with a health cost as it sacrifices the thermal benefits of clustering^45^. This has been demonstrated, for example, in free-ranging kudu, where sick individuals sought warm environments to support the febrile response^20^. Interestingly, we observed one healthy female that consistently approached sick-like isolated females, which she had been housed with for months, and hung near them. While this is only a single anecdote, it is the exact opposite of what we would expect if healthy individuals were avoidant in order to protect themselves from catching a disease. Additionally, it indicates the possibility of a role of social history or demographics in moderating behavior between healthy and sick individuals, such as in vampire bats^32^.

One explanation for self-isolation is that the behavior benefits group members, providing an indirect benefit to the sick individual. Shakhar and Shakhar^5^ propose sickness behavior serves to broadly reduce disease transmission, and should be present in response to highly virulent and highly transmissible diseases where isolation does little to change the outcome on an individual level, but has high potential benefits if it prevents disease from spreading to kin. They further state that such behaviors should be emphasized in species living in close proximity, such as Egyptian fruit bats, where chances for transmission are very high. Isolation for kin benefit has only been documented in eusocial species, where individuals are highly related to group members. Previous analysis suggests that roost mates of Egyptian fruit bats are not more related to one another than would be expected by chance^46^, making kin selection an unlikely evolutionary driver.

Alternatively, removal from the cluster may directly benefit the infected individual in multiple ways. Hanging in a less insulated location may keep an individual’s fever from becoming dangerously high. Additionally, bats within clusters often squabble and push one another around. By not being in the cluster, sick individuals would not have to expend energy on interindividual conflict or maintaining cluster position. Finally, self-isolation may prevent an already sick individual from catching an additional illness, as multiple concurrent illnesses are far more deadly than lone infections^47^. While most likely driven by benefits to the individual, this behavior may serve to reduce transmission, and benefit the group as a by-product, meaning kin selection may not be required to bring about such group-benefitting behaviors. This emphasizes the importance of considering the benefits to the self and others together in the evolution of sickness behavior, rather than assuming they are competing processes. Like humans reducing contacts when sick^48^ or social distancing efforts made in 2020 to slow the spread of Covid-19^49^, increasing space between individuals may reduce infection rates, particularly in densely populated groups such as the ones Egyptian fruit bats live in.

This study is the first to record changes in foraging of sick-like bats in the field. All sick-like individuals in the open colony failed to exit to forage for at least two nights. This was clearly not exclusively a result of food supplement as these bats always flew out to forage prior to the treatment and as they consumed little food during the treatment, similar to the decrease in food consumption observed in the closed colony. While we cannot exclude the possibility that fully wild bats may engage in short flights while sick, the clear pattern of well-known sickness behaviors, such as anorexia and lethargy, due to the acute phase response of the immune system, documented here makes this highly unlikely. These findings demonstrate the lack of food consumption widely observed in laboratory and livestock settings^1,50–56^ also occurs in free-ranging animals, supporting the highly conserved presence of anorexia in sick animals. As our bats in the open colony exhibit foraging behavior very similar to that in other wild bats^57^ we predict that this behavior is general for other bat species as well. In the wild, foraging can take considerable effort and exposes animals to predation risk, particularly for individuals that are not in good condition. Thus, staying in place and not traveling to forage benefits sick animals by conserving energy and increasing safety. Anorexia is also thought to support the immune response to bacteria^58,59^ and laboratory studies in mice have demonstrated increased survival in individuals with restricted food intake, either prior to^59^ or during^60^ infection. Finally, it may also reduce disease transmission by decreasing the number of contacts between sick individuals and shared food sources^5^.

The acute phase response has been examined in six species of bats, revealing much variability. Some show clear leukocytosis (*Chaerephon plicatus*^61^, *D. rotundus*^30^) but in others this response is not clear (*A. lituratus*^27^), totally absent (*M. molossus*^28^), or present in some scenarios (*C. perspicillata*^29^) but not in others (*C. perspicillata*^26,31^). We did not find leukocytosis. Normal, healthy captive *R. aegyptiacus* display an extremely wide range in leukocyte count (see Zims^62^), making it difficult to follow leukocytosis even if it did occur. We believe that a significant leukocytosis in the challenged group might have been masked by a stress leukogram (also known as NLR, see: Davis et al^63^) that was caused by the handling of bats from both groups, reflected in elevation of neutrophil and decrease in lymphocyte counts in the blood sample before the challenge. Stress leukogram was found to maintain the higher ratio in the challenged group, similar to previous findings in *D. rotundus*^30^ and food deprived *C. perspicillata*^26^ in reaction to LPS challenge.

We observed a clear elevation in haptoglobin, an acute phase protein of the innate immune response, following the LPS challenge. Haptoglobin has only been measured in a few bats, which showed a clear elevation in response to fungal^34^ and human stress^38^. In both free and captive bats, haptoglobin continuously increased during the entire period of the study, as also observed in other mammals^64^. It is interesting to see that even at baseline some individuals’ haptoglobin level was not zero, perhaps due to the stress of handling. The free ranging colony’s higher baseline levels could have been due to the younger age of these bats, or to their nightly potential exposure to bacterial pathogens outdoors.

Lysozyme levels were higher following LPS injection in the experimental group in the closed colony but not in the open colony. This finding has several potential explanations. First, the open colony was examined only pre-challenge and 48 hours post-challenge, which was later than the peak in lysozyme levels at 24 hours post-challenge in the closed colony, thus the lysozyme levels may have decreased by that point. Second, there may be an age difference between adults and adolescents in the lysozyme level, as was found in bison^65^. Third, the relatively small number of bats examined in the open colony and the high variation between individuals may have obscured any potential effect.

## Conclusion

Here we clearly demonstrate that in addition to following many aspects of classical mammalian sickness physiology and behavior when faced with a bacterial-type immune challenge, Egyptian fruit bats exhibit self-imposed social distancing. This is the first time social self-isolation has been recorded in a mammal, rather than isolation through avoidance by conspecifics or as a by-product of lethargy. Such isolation behavior stands in stark contrast to normal behavior in this species, reflecting a temporary shift in needs to prioritize survival.

Anorexia, a well-documented aspect of sickness behaviors, is here, for the first time, documented in the context of staying in the sleeping shelter, as expected based on our current understanding of the role of anorexia during bacterial infection. The bats’ self-isolation together with their staying in the colony suggests how sickness behavior can reduce the transmission of pathogens both inside and outside the colony, including reducing the probability of inter-species spillover events. This supports previous findings that spill-over events are primarily caused by human disturbance^67^, emphasizing the importance of leaving critical habitat features, such as roosts, alone for reducing the likelihood of future events like the Covid-19 pandemic.

Immunologically, the response to bacterial infections displayed both similarities and differences to other bat species and differed from how the same species responds to viral infections^66^. Immune-challenged individuals did not have clear leukocytosis but had a significantly higher neutrophil lymphocyte ratio (NLR). Molecularly, we are the first to measure elevation in haptoglobin and lysozyme in bats during a bacterial immune challenge. To our knowledge, this study is the first time an LPS challenge was followed for 48 hours post challenge in bats or other mammals and it is one of the few studies which uses a long lived, social species as a model. Our unique findings in this species clearly indicate the importance of both life history features and social patterns on responses to adversity, such as disease.

The acute phase response in *R. aegyptiacus* across multiple behavioral and physiological measures clearly prioritizes bacterial eradication and host recovery over other short- and long-term selective pressures, protecting both the sick individual and their many group members. Our findings provide insight into the multiple ways that highly social, long lived mammals have evolved to adjust their behavior during illness to mitigate both the stress imposed upon sick animals living in dense groups and to reduce the high pathogen transmission risk inherent to such groups.

## Methods

### 1. Experimental animals

#### a. Closed Colony

Nineteen adult male bats were captured from a cave roosting colony in Herzliya, Israel on the 2nd of December 2018. They were checked for ectoparasites and treated topically with carbaryl powder to reduce parasitic load (Opigal, Abik, *88000029*). The males were housed together in the experimental room (245 cm × 200 cm × 210cm) for two weeks for acclimation to captivity and handling. Females (n=18) were brought from a captive colony consisting of mothers and pups in order to avoid pregnant females in the experiment. An additional room (4m x 2m x 2.9m) was used as a temporary housing space as needed for individuals not actively in the experiment.

The temperature in the experimental room was maintained at 25 □ using a wall mounted heater/AC unit. Natural dark-light cycle was enabled by some light penetration from the outside during the day. A variety of diced fruit (banana, apple, and melon) of 150 grams per individual was provided daily. All individuals were marked with a unique symbol using hair bleach on their heads and backs for identification purposes.

#### b. Open Colony

A total of five individuals (three females and two males) were selected from the open colony located at Tel Aviv University^39^ for this experiment. These were healthy young adults aged approximately seven months that had been observed to regularly travel away from the colony and return by dawn. All bats had access to the outdoors and were provided with diced fruit. Individuals were marked with a unique symbol using hair bleach for identification on their head.

Animal care and experimental procedures were approved by Tel Aviv University IACUC under approval form ID 04-19-002.

### 2. Immune challenge

Closed colony individuals were randomly assigned to the control group, treatment group, or the cluster group, which was defined as individuals not measured in the experiment, but housed in the room to provide a richer social environment for the experimental bats. Open colony individuals served as their own controls.

Individuals in the treatment group were injected subcutaneously with *Escherichia coli* O111:B4 lipopolysaccharides (LPS, Sigma Alrdich, L2630) at a concentration of 4 mg/kg b.w., (0.577±0.144mg) diluted in sterile phosphate buffered saline (PBS, Sigma-Aldrich, P5493), while control animals received the same amount of PBS. Dosage was determined in a preliminary experiment comparing the clinical outcome (body temperature elevation and visible lethargy and joint swelling) of injecting 2 mg LPS /kg b.w. and 4 mg LPS /kg b.w.. In order to be more cautious, considering the age of the individuals for the open colony, a reduced dosage of 2 mg\kg b.w. (0.204±0.031mg) was confirmed as effective in achieving a similar outcome and was used in all these individuals.

### 3. Experimental design and data collection

#### a. Closed Colony

The data collection on the closed colony was conducted in two rounds. Males (M) and females (F) were placed together three days before each round in order to minimize possible pregnancies while creating a normal mixed-sex colony. The first round contained five (3F, 2M) challenged, five (2F, 3M) control, and 13 (5F, 8M) cluster individuals. The second round contained 6 challenged (3F, 3M; 1F deceased), five (4F, 1M) control, and six (2F, 4M) cluster individuals. Six (1F, 5M) of the non-challenged bats from the first round were retained for the second round. Of these, two bats that were not measured (part of the cluster) in the first round were selected to participate in the second round to avoid any possible influence on physiological measurements, the other four (1F, 3M) were part of the cluster. One female died during the course of the experiment, and these data were excluded from the analysis.

For both rounds, the bats were handled five times, consisting of four time points for data collection (pre-injection, 12, 24, and 48 hour post-injection) and the injection time. In order to avoid confounding effects from circadian changes in body temperature, as known for this species^40^ we challenged each one of the rounds in a different part of the day; the first group at night, and the second in the morning. For the first experiment, these times were as follows: pre-injection between 11:30-14:00, injection with LPS or PBS at 21:00-22:30, 12h between 10:00-12:00, 24h post injection at 22:00-00:00, and 48h post injection at 22:00-00:00. Handling times for the second round were as follows: pre-injection at 16:00-20:00, injection at 10:15-12:45,12 h post injection at 22:00-00:00, 24 h post injection at 10:30-13:00, 48 h post injection at 10:30-13:00.

During handling, challenged and control bats were removed from the colony to measure body weights and collect blood samples. All individuals were offered mango juice immediately following handling.

For both rounds, approximately 250 μl of blood was obtained via venipuncture with microtainer tubes treated with K2EDTA (BD reference 365,974) from the antebrachial or the wing vein using different locations for each sampling. Blood was kept cold and later centrifuged at 8000 RPM for three minutes for complete separation between the packed cells and the plasma. Plasma was collected to a new Eppendorf tube and stored at −80°C until further analysis. Additionally, at least three blood smears were made for each individual each time blood was collected. Immediately after blood was drawn, blood smears were stained using a May-Grunwald eosin methylene blue solution (Merck-Millipore catalog number 10142401251730), as described in Schneeberger^68^.

During the pre-injection handling small biologger devices (Vesper, ASD inc.) coated in Parafilm (Heathrow Scientific) and duct tape were glued to the back of the bat using Perma-Type Surgical Cement (AC) to three challenged bats and to two control bats in round 1 and all experimental bats in round 2. These devices were gently removed during the 48h post-injection handling. In both rounds, biologgers were set to measure skin body temperature at 2-minute intervals through a sensor placed on the back over closely trimmed fur. In round 2, the biologgers additionally measured acceleration at 50 times per second. Infrared video cameras were used to continuously monitor the bowl of provided food to track food consumption in both rounds. In the second round, additional cameras were added to monitor the ceiling to track social isolation.

#### b. Open Colony

Data collection on free-ranging individuals began on the 28th of February and ended on the 12th of April. Each bat was fitted with a datalogger attached to a collar containing an RFID tag. All individuals had GPS recorded; three individuals also had acceleration recorded. Video and RFID were used to record entrances to and exits from the open colony. Provided food and the ceiling of the colony room were continuously monitored using infrared cameras in order to track food consumption and social isolation, respectively. Blood was collected in the same manner described above only twice for the young adults, immediately prior to the challenge and 48 hours post-challenge.

For two individuals, a PBS control was conducted multiple days after their LPS trial, the other three had the saline control prior to injection with LPS. Baseline behavioral data covered two days of social distance records and four days of GPS tracking for each individual.

### 4. Video analysis

#### a. Food consumption

##### i. Closed

Infrared video of the provided food in the closed colony was processed by two independent coders with greater than 80% reliability with each other and KRM on 20% of the video data. All occurrences of accessing the food bowl were recorded with the bat’s ID, number of pieces of fruit consumed, and whether those pieces were banana or not.

##### ii. Open

Two independent coders each responsible for a subset of the individuals processed video footage of the provided food. Their work was checked by KRM. Video from the dates an individual bat did not exit were monitored to record the number of pieces the focal bat consumed when they stayed inside the colony room.

#### b. Social Isolation

Videos of the colony rooms were manually processed to determine isolation levels. For the closed colony, each individual’s location on screen was recorded at 10 minute intervals and used to calculate their distance from their nearest neighbor. Individuals within the cluster were recorded as having a standard inter-individual difference based on multiple measurements of distances between bats in the cluster, as it was not possible to always identify individuals while they were clustered. For the open colony, each individual’s isolation level was recorded in 30-minute intervals on a 0-3 scale. Isolation levels were defined as being within a cluster, in contact with another individual, within two body widths of another individual (approximately 16cm), or greater than two body widths from any other individual.

### 5. Laboratory analyses

#### a. White Blood cell counts

To assess changes in leukocyte counts, four blood smears were analyzed from each bleeding session (prior, 12h, 24h and 48h post injection). Immediately after blood was drawn, blood smears were stained using a May-Grunwald eosin methylene blue solution (Merck-Millipore catalog number 10142401251730). For every smear, leukocytes, (categorized as neutrophils, lymphocytes, monocytes, and eosinophils) were counted manually up to 100 cells per slide at ×□400 magnifications. The mean of each leukocyte type counted in three smears was calculated as an estimate of each leukocyte cell type count per individual for each of the four time points. The neutrophil lymphocyte ratio (NLR) was calculated per individual for each of the four time points by dividing the neutrophil count by the lymphocyte count.

#### b. Haptoglobin concentration

To measure the concentration of haptoglobin, the standard procedure of the commercial kit “PHASE”TM Haptoglobin Assay (Tridelta, Maynooth, Ireland) was followed, which was already used in other bat species^34,38^. Briefly, after diluting the plasma samples (1:2) with PBS, haemoglobin was added. Haptoglobin binds to haemoglobin and maintains its peroxidase activity at a low pH. The measured peroxidase activity of haemoglobin is directly proportional to the amount of haptoglobin in the sample. Haptoglobin concentrations were calculated according to the standard curve on each plate and were expressed as mg/ml.

#### c. Lysozyme concentration

To measure lysozyme concentration, we used the lysoplate assay, which was adapted to low sample volumes^38,69^. Briefly, we prepared 1% noble agar (Sigma Aldrich) with PBS at pH = 6.3 and we added the required amount of lysozyme-sensitive bacteria Micrococcus lysodeikticus ATCC #4698 (Sigma Aldrich) to reach a bacterial concentration of 50 mg/100 ml in the agar. We poured the agar in Petri dishes on a pre-heated surface (50°C) and horizontally leveled with a spirit level to avoid quick and uneven cooling of the medium. After solidification, we inoculated 1.5 µl of plasma in test holes (1.7 mm in diameter). We used standard dilutions of hen egg white lysozyme (0.5, 0.8, 1, 2, 4, 8, 10, 20, and 40 µg/ml; Sigma-Aldrich) to prepare a standard curve in each plate. We incubated the plates at 37°C for 20 hours. During the incubation, a zone of clearing developed in the area of the gel surrounding the sample inoculation site as a result of bacterial lysis. The diameters of the cleared zones are proportional to the log of the lysozyme concentration. We photographed each plate in a photobox (Imaging system; peqlab) with a ruler next to it as a reference scale. We measured the diameter of the cleared zone digitally three times using the software ImageJ (version 1.48, http://imagej.nih.gov/ij/) and we converted the mean value on a semilogarithmic plot into hen egg lysozyme equivalents (HEL equivalents, expressed in μg/ml) according to the standard curve^70^.

### 6. Statistical analysis

#### a. Evidence of Illness

##### i. Temperature

The first five temperature records of each two-hour increment were averaged to provide bi-hourly samples from individuals in the closed colony. To determine differences in temperature immediately following LPS injection, a Mixed ANOVA was used to compare challenged (n=5) and control (n=6) groups bi-hourly from six hours prior to injection to eight hours post injection. To visualize temperature change across the full experiment, all temperature measures were smoothed using a moving average.

##### ii. Acceleration

Accelerometry measurements from the second round of data collection in the closed colony were smoothed using a moving average and filtered using a Butterworth filter, then values from all three axes were combined using a norm. For each combination of day/night and pre/post injection, each individual’s proportion of time above 30% of the maximum acceleration value for that individual was calculated to indicate amount of non-minor movement for each time period in the study. The proportion of samples above the threshold was compared between challenged (n=4) and control (n=4) individuals pre- and post-injection and during day and night using a mixed-measures GLMM. An insufficient number of individuals from the open colony had complete acceleration recordings for analysis.

##### iii. Food Consumption

Food consumption in the closed colony was compared between challenged (n=10) and control (n=10) individuals across nights using a mixed ANOVA.

##### iv. Weight

Weights measured from the closed colony were compared between challenged (n=10) and control (n=10) groups across the four time periods using a mixed ANOVA. For the open (n=5) colony, weights pre- and 48 hours post-injection were compared using a repeated measures t-test.

#### b. Sociality/isolation

Records of distance from each individual to their nearest neighbor from both the closed (Challenged n=5; Control n=5) and open (n=5) colony were analyzed with a generalized linear mixed effects model.

#### c. Foraging Behavior

##### i. Proportion leaving

We used a Fisher’s exact test on two control nights and two nights immediately following treatment to determine the effects of LPS on exiting behavior in the open (n=5) colony.

##### ii. GPS

Total distance traveled was calculated from GPS records of 4 control nights and the 4 nights immediately following injection. Distances were converted to Z-scores for each individual (n=5), and then compared between control and treatment conditions across nights using a general linear model.

##### iii. Food Consumed

Food consumption on the first two nights an individual (n=5) did not exit the colony room was compared to the number of food pieces each individual was expected to eat based on body weight and average weight of fruit pieces provided using a repeated measures ANOVA and Tukey’s post-hoc test.

#### d. Immunological measures

White Blood cell count, NLR, Haptoglobin, and Lysozyme were all compared using the same tests. For the closed colony, comparison between control (n=10) and challenged (n=10) groups across the four sampling periods was done using a Mixed ANOVA. For the open (n=5) colony, comparison between pre- and 48 hours post-injection was done using a repeated measures t-test.

## Acknowledgments

We are grateful to Katja Pohle for her crucial help with the laboratory analysis. We thank Linoy Alkalay, Eidan Loushi, Liad Pinchasi, and Nofar Gergrood for their contributions to processing many hours of video, Sharon Krelenstein and Noam Golan for counting white blood cells, and Reut Assa for bat handling assistance. We appreciate the technical assistance of Aya Goldstein with the Vespers and Ofri Eitan with the cameras. Thanks to Zohar Tal for helping rear the pups used in the open colony portion of the experiment. KRM was supported by the Zuckerman STEM Leadership Program. GÁC was supported by funds from the Leibniz Institute for Zoo and Wildlife Research, Berlin. LGHM was funded by Consejo Nacional de Ciencia y Tecnología (#237774). This research was supported by the Israel science foundation, grant number 677/17.

## Ethical approval

All applicable institutional and/or national guidelines for the care and use of animals were followed. Animal care and experimental procedures were approved by Tel Aviv University IACUC under approval form ID 04-19-002.

## Author contributions

MW, GÁC, YY, LGHM conceived of and designed the overall study

KRM conceived of and designed the social behavior component of the study

LH conceived of, designed, and implemented the GPS tracking component of the study

MW, KRM, VBSR conducted closed colony data collection

MW, LH, KRM conducted open colony data collection

MW, GÁC conducted laboratory analyses on immunological measures

KRM conducted statistical analyses

KRM, MW drafted the manuscript (with edits from YY and GÁC)

All authors contributed to and approved of the final manuscript

## Supplemental figures

**Supplemental Figure 1.**
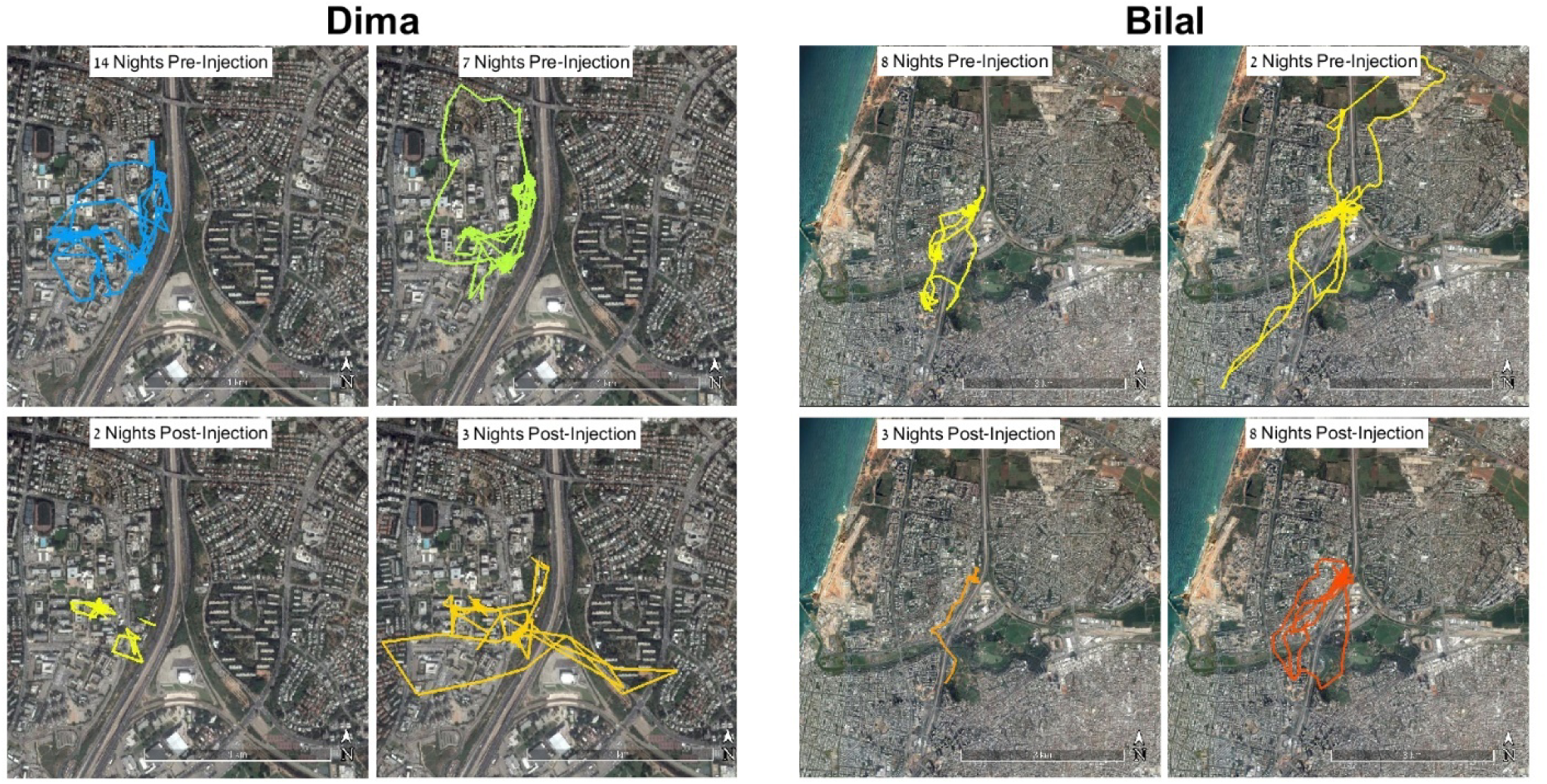
GPS tracks for two individuals. Shown are two nights prior to LPS-challenge were selected to demonstrate normal nightly foraging behavior (top), the first night exiting after exiting behavior ceased in response to the LPS challenge (bottom left of each group of four images), and the first night each individual returned to a distance similar to that covered prior to challenge (bottom right of each group of four images).

**Supplemental Figure 2.**
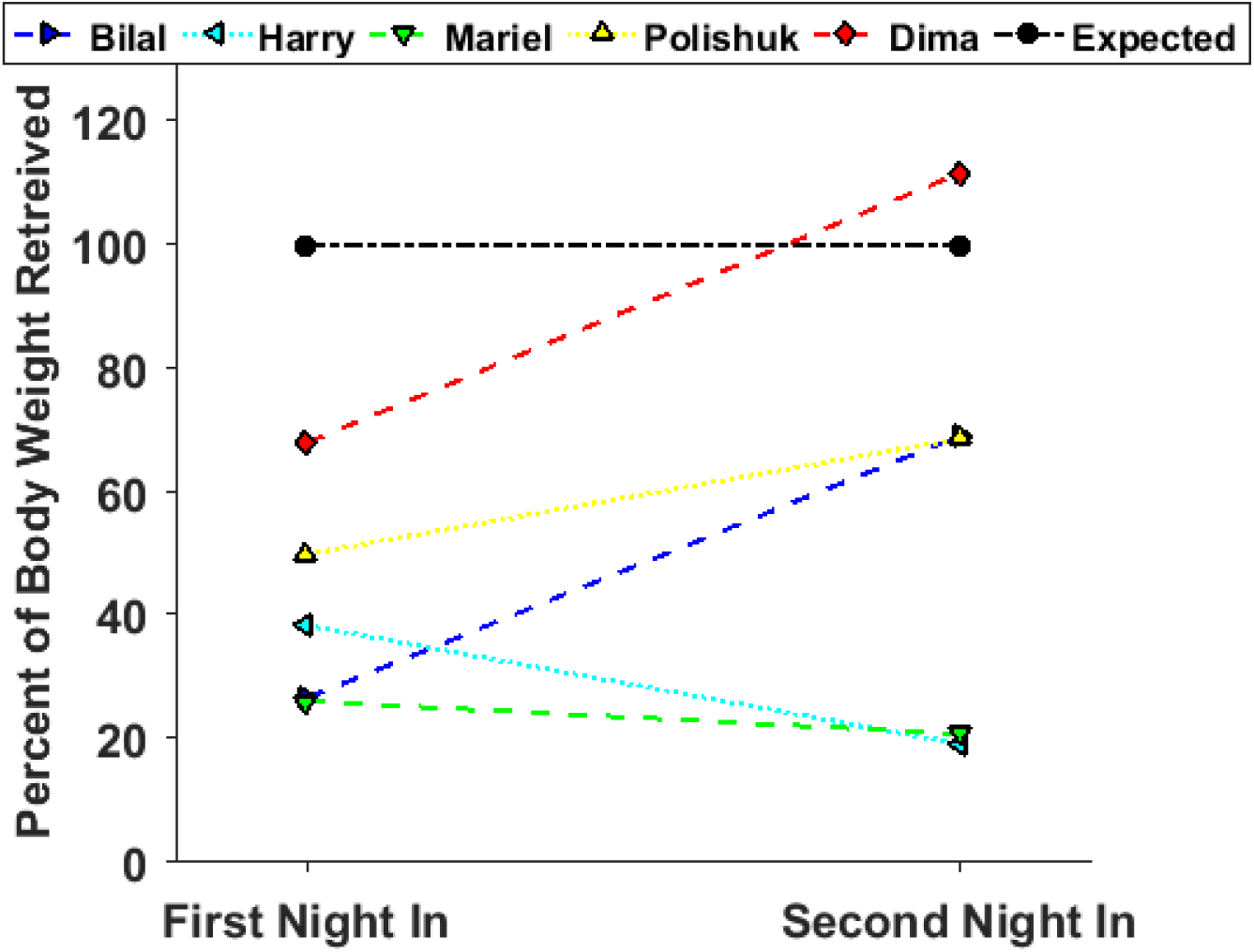
Individuals ate less than expected (shown as a black line) during their first two nights staying inside the colony

