## Supplementary figures and images for "Sick Bats Stay Home Alone: Social distancing during the acute phase response in Egyptian fruit bats (*Rousettus aegyptiacus*)"

### Supplementary figure 1

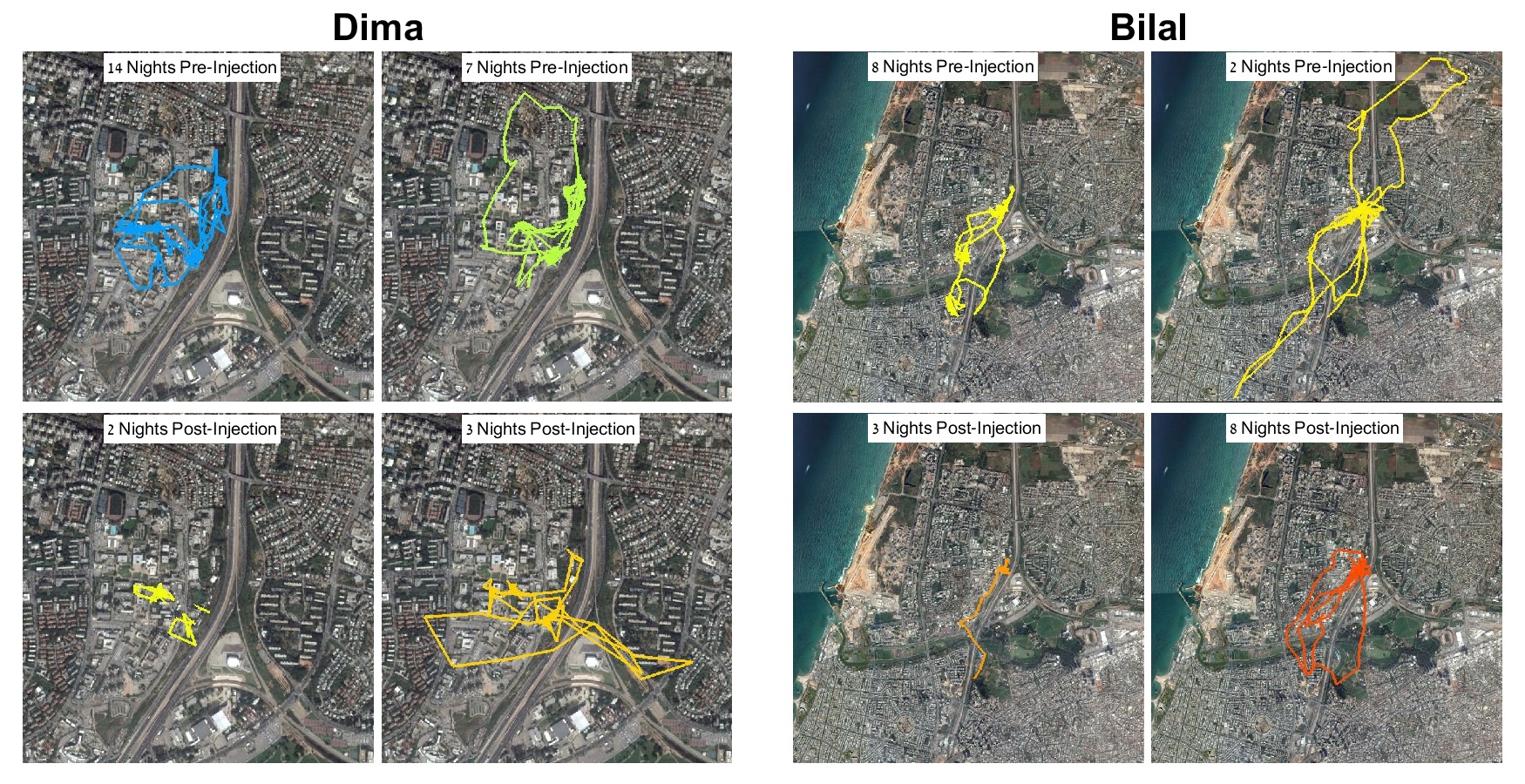

### Supplementary figure 2

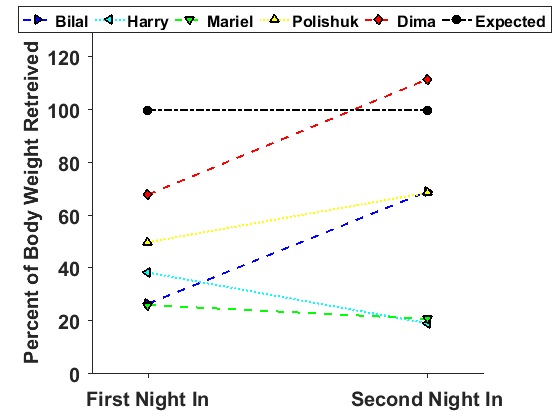
